# Heating tolerance of ectotherms is explained by temperature’s non-linear influence on biological rates

**DOI:** 10.1101/2022.12.06.519315

**Authors:** Jacinta D. Kong, Jean-Francois Arnoldi, Andrew L. Jackson, Amanda E. Bates, Simon A. Morley, James A. Smith, Nicholas L. Payne

## Abstract

The capacity of ectotherms to adjust their thermal tolerance limits through evolution or acclimation seems relatively modest and highly variable, and we lack satisfying explanations for both findings given a limited understanding of what ultimately determines an organism’s thermal tolerance. Here, we test if the amount of heating an ectotherm tolerates throughout a heating event until organismal failure scales with temperature’s non-linear influence on biological rates. To account for the non-linear influence of temperature on biological rates on heating tolerance, we rescaled the duration of heating events of 316 ectothermic taxa acclimated to different temperatures and describe the biological rate-corrected heating duration. This rescaling reveals that the capacity of an organism to resist a heating event is in fact remarkably constant across any acclimation temperature, enabling high-precision estimates of how organismal thermal tolerance limits vary under different thermal regimes. We also find that faster heating consistently reduces biological rate-corrected heating durations, which helps further explain why thermal tolerance limits seem so variable on absolute temperature scales. Existing paradigms are that heating tolerances and thermal tolerance limits reflect incomplete metabolic compensatory responses, are constrained by evolutionary conservatism, or index failure of systems such as membrane function; our data provide a different perspective and show that an organism’s thermal tolerance emerges from the interaction between the non-linear thermal dependence of biological rates and heating durations, which is an approximately-fixed property of a species.

## Introduction

The thermal tolerance limits of organisms are of increasing focus for understanding how rising temperatures and extreme heat events shape ecology and evolution (Sinclair *et al*. 2016), and for predicting organismal and community resilience to climate warming (Morley *et al*. 2019b). Recent comparative analyses indicate ectotherms have some capacity to increase their thermal tolerance limits through short-term acclimation or adaptation (Rohr *et al*. 2018; Salinas *et al*. 2019), but upper thermal tolerance limits only vary marginally both within and across species, especially when compared to lower thermal tolerance limits (Araújo *et al*. 2013; Gunderson & Stillman 2015). This apparent low flexibility of upper thermal tolerance limits across lineages from different climates, latitudes or altitudes, as well as in acclimation experiments, raises concerns about the ability of ectotherms to mediate physiologically the impact of future climate warming (Hoffmann, Chown & Clusella-Trullas 2013; Gunderson, Dillon & Stillman 2017). As flexibility in physiological responses is undoubtedly important for the persistence of species in the face of heat events unfolding over different time scales (Donelson *et al*. 2019; Morley *et al*. 2019a; Bates & Morley 2020; Fredston-Hermann *et al*. 2020), there is a need to improve our understanding of the factors that regulate organismal thermal tolerance limits (Chung & Schulte 2020; Terblanche & Hoffmann 2020; van Heerwaarden & Kellermann 2020).

Upper thermal tolerance limit, T_c_, is defined in biological terms as the temperature at which failure occurs when an organism is heated. The physiological and biochemical mechanisms that cause failure at T_c_ are debated, with leading explanations involving changes in membrane fluidity or protein function that respond to high temperature *per se* (Bowler 2018). However, an increasing body of work shows that T_c_ also depends on the rate of heating (slower heating decreases T_c_)(Kingsolver & Umbanhowar 2018) and that the duration an organism can withstand exposure to a static high temperature varies systematically with the temperature of exposure (faster onset of failure at higher temperatures)(Santos, Castañeda & Rezende 2011; Rezende *et al*. 2014). Furthermore, there is increasing evidence that the onset of failure might be linked with biological rates through processes such as the expenditure of metabolites or injury accumulation (Santos, Castañeda & Rezende 2011; Jørgensen *et al*. 2021). Taken together, the dependence of thermal tolerance limits on both temperature and time implies failure at high temperature must be mechanistically related to biological rates. That is, whatever the ultimate physiological mechanisms that cause organismal collapse at T_c_ (e.g. membrane fluidity)(Pörtner 2002), data show that the onset of collapse is dependent on the duration of heating and therefore biological rate dynamics, not just temperature *per se*.

Increasing an organism’s body temperature implies heating (a rise in temperature) applied for a given duration of time. As temperature increases, biological rates such as metabolic rate increase exponentially following a consistent Arrhenius form ∼ e^−0.6/kT^ (the Universal Temperature Dependence), where k is Boltzmann’s constant and T is temperature (Brown *et al*. 2004). If the capacity of an organism to tolerate a heating event is determined by the cumulative expenditure of some currency (e.g. energy, metabolites or water)(Santos, Castañeda & Rezende 2011) or accumulation of cellular damage up to a limit (Jørgensen *et al*. 2021), then the onset of organism failure at T_c_ (or after some duration of exposure to a static high temperature) will be linked to biological rate dynamics because these processes (energy expenditure, damage accumulation) manifest as rates. It follows that the heating tolerance of an organism may be directly determined by the total magnitude of physiological currency (energy, metabolites, or damage) that is transformed during heating, at a rate that is temperature-dependent following the Universal Temperature Dependence. If so, quantifying this currency could help explain how and why thermal tolerance limits vary, and also improve our ability to predict thermal tolerance limits under different conditions. While the scaling of biological rates with temperature (dictated by reaction kinetics) and the failure at T_c_ are often considered independent phenomena, an increasing suite of empirical data implies they are not. For example, by measuring how exposure temperature scales with time-to-death (commonly the *z* parameter), other studies have phenomenologically quantified how ‘injury accumulation rate’ exponentially increases with temperature and used those observed relationships to successfully predict thermal limits under different conditions (Jørgensen, Malte & Overgaard 2019; Jørgensen *et al*. 2021). However, those studies rely on measuring observed dependencies of temperature on thermal limits to formulate predictions, and do not explicitly consider the Universal Temperature Dependence of biological rates. Santos and colleagues (2011) incorporated metabolic costs into simulations of how water depletion impacts *Drosophila* thermal limits, but whether invoking the Universal Temperature Dependence helps explain thermal tolerance limits in ectotherms in general is unclear.

In this study, we investigate the role of temperature, time, and non-linear biological rates in generating variation in upper thermal tolerance limits of diverse ectotherm species. To do this, we consider a large dataset that reports the upper thermal limits of 316 species from 7 Phyla when they are heated, at a consistent rate, from two different acclimation temperatures (T_a_) until the temperature at physiological failure (T_c_). Using an integration approach and scaling biological rates ∼ e^−0.6/kT^ (the Universal Temperature Dependence)(Brown *et al*. 2004), we computed the effective elapsed duration of the heating event that accounts for the non-linear scaling of biological rates with temperature (Δt_r_). We find that this biological-rate-corrected duration of a heating event, which we call ‘rescaled heating duration’ Δt_r_, is remarkably constant for an organism acclimated to very different temperatures after the Universal Temperature Dependence is accounted for. These rescaled durations are consistent to the extent that we consider Δt_r_ to be an approximately fixed property of a species. We additionally find that heating rate strongly and consistently influences Δt_r_. Our study offers a complementary perspective to recent studies linking temperatures to times or rate dynamics in order to understand what regulates thermal tolerance limits (Santos, Castañeda & Rezende 2011; Rezende *et al*. 2014; Kingsolver & Umbanhowar 2018; Jørgensen *et al*. 2021), and reveals a strong quantitative link between the familiar scaling of biological rates underpinning the metabolic theory of ecology (Brown *et al*. 2004) and the onset of organismal collapse. The data also raises questions about the true capacity of ectotherms to adjust their thermal tolerance limits.

## Materials and Methods

### Heating tolerance and heating duration

We defined the range of temperatures spanning a heating event (e.g. a heating assay) as heating tolerance, H (Fig. 1A),

**Figure 1.**
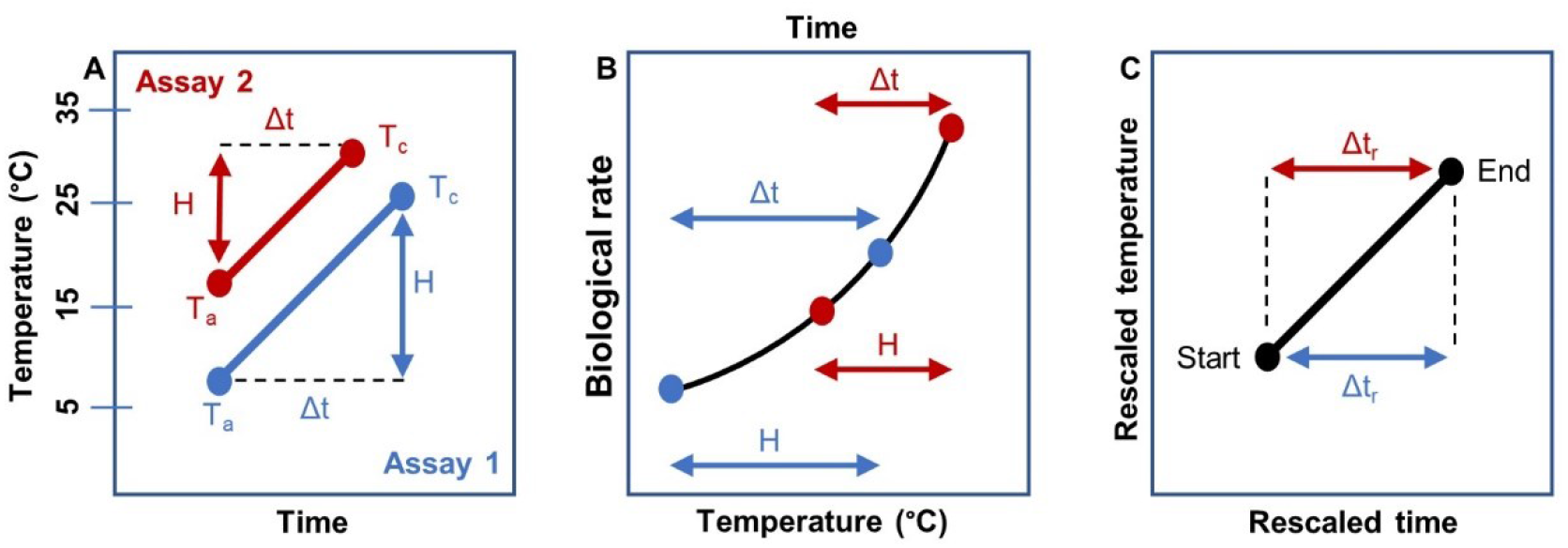
A) Heating tolerance (H = upper thermal tolerance limit T_c_ – acclimation temperature T_a_) of a thermal tolerance assay at a lower acclimation temperature (Assay 1; blue) is larger than the heating tolerance at a higher acclimation temperature (Assay 2; red) with the same heating rate. B) Under the same heating rate, organisms attain upper thermal tolerance limits τ_c_ faster (Δt) at higher acclimation temperatures (red) than lower acclimation temperatures (blue). C) Elapsed durations rescaled for the Universal Temperature Dependence (Δt_r_) are the same for both acclimation temperatures on normalised temperature scales.

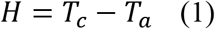

where T_a_ is acclimation temperature (°C) and T_c_ is upper thermal tolerance limit (°C). We reanalysed a published dataset to calculate heating tolerances (H) of 316 ectotherms from 7 Phyla where each species had heating tolerance measured from two different acclimation temperatures, which were on average ∼ 10°C apart (Morley *et al*. 2018; Morley *et al*. 2019b) (Supplementary Material). As a heating event with a constant heating rate involves an increase in time and temperature, we defined an elapsed duration of the heating event corresponding to H (Fig. 1A),

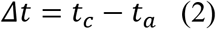

The elapsed duration of the heating assay, Δt, begins at the start of the ramping assay (t_a_; e.g., 0 minutes) and ends at the onset of T_c_ that signals the end of the ramping assay (t_c_). Thus, these times correspond to a temperature in degree Celsius (i.e., t_a_ at T_a_ and t_c_ at T_c_).

### Rescaling heating durations for non-linear temperature-biological rate relationships

To quantify the relative quantity of biological processes accumulating over time throughout a heating event from T_a_ to T_c_ and rescale heating tolerances, we used the Universal Temperature Dependence of metabolism (Brown *et al*. 2004) as a proxy for the non-linear temperature-dependence of biological rates (Fig. 1B). The rescaled heating duration (Δt_r_) is corrected for non-linear biological rates and acclimation temperatures. We consider Δt_r_ a relative rather than absolute quantity to allow for more general interpretations of the currencies that it might represent (such as energy [ATP], metabolites, water, or cumulative damage) and to account for the non-linear scaling of biological rates with temperature.

To simplify the rescaling of temperature-time dynamics according to the Universal Temperature Dependence, we centred temperatures in degree Celsius (T) to a reference temperature (0°C; the normalisation constant T_K_ = 273.15) and used this dimensionless normalised temperature scale (τ = T / T_K_; no units) to approximate, up to a multiplicative constant, the Universal Temperature Dependence as

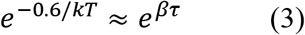

where *β* = 0.6/kT_K_ is a dimensionless constant constructed from the organism-level activation energy, Boltzmann’s constant k, and the normalisation constant T_K_. *e*^*βτ*^ is the dimensionless biological rate-temperature relationship that is an equivalent expression of the Universal Temperature Dependence on a normalised temperature scale (τ) (Supplementary Material). The value of activation energy used to define *β* is not fitted to our data and changing this value (E = 0.6eV) within realistic bounds does not meaningfully change our results (Supplementary Material, Fig. S1). Normalised temperatures can be rescaled back into degree Celsius by T = τ ×T_K_.

The relationship between temperature and time shared via biological rates on Celsius temperature scales (T) can be equivalently expressed on normalised temperature scales (τ) (Fig. 1B). Normalised heating tolerance (Δ*τ* = τ_c_ - τ_a_) are equivalent to heating tolerance (H) on the normalised temperature scale with τ_c_ and τ_a_ corresponding to T_c_ and T_a_ respectively. Δ*τ* can be converted to H in degrees Celsius by H = Δ*τ*×T_K_, i.e, Δτ = τ_c_ - τ_a_ = T_c_/T_K_ - T_a_/T_K_ = H/T_K_. Both Δ*τ* and H have the same corresponding time elapsed during the heating assay (Δt).

The dimensionless biological rate-temperature relationship (*e*^*βτ*^) can be expressed as a function of normalised temperature changes over time, *e*^*βτ*(*t*)^, where τ is defined as normalised temperature at time t. Hence, we can express the time elapsed between the start of the ramping assay (t_a_) and the end of the ramping assay (t_c_) as an integral of the dimensionless biological rate-temperature relationship *e*^*βτ*(*t*)^ over time and account for the non-linear change in biological rates with temperature over the time-course of the assay (Eqn. 4).

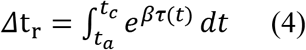

Δt_r_ is the biological rate-corrected elapsed duration of the ramping assay (rescaled heating duration; Fig. 1C), that is, the effective duration of the heating event that determines the total biological rate output of the organism during the heating event (e.g., total oxygen consumption as a proxy for metabolic rate). Δt_r_ is the rescaled equivalent of elapsed duration of the heating event (Δt). Subscript _r_ denotes biological rate-corrected variables (Table S1).

### Rescaling heating tolerance for non-linear temperature-biological rate relationships and heating rate

If heating rate λ (expressed on the normalised temperature scale; λ = heating rate (°C time^-1^) / T_K_; units: time^-1^) is constant during the heating assay, time and temperature become equivalent variables (since dτ = λdt) and the integral in Eqn. 4 can be simplified as

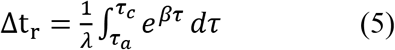

Eqn. 5 describes the cumulative quantity of a biological process transformed in response to heating over the elapsed duration of a heating event that includes the heating rate of a ramping assay. Integration of Eqn. 5 gives an expression of rescaled heating duration (Δt_r_) on the normalised temperature scale (τ), allowing for Δt_r_ to be calculated for any heating tolerance (H or Δτ) (Eqn. 6). Eqn. 6 can be simplified by defining the function *g*_*β*_*(x)*.

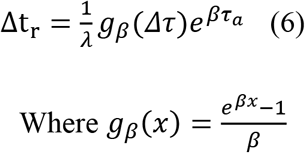

The inverse of Eqn. 6 converts rescaled heating duration (Δt_r_) into normalised heating tolerance (Δτ) (Eqn. 7).

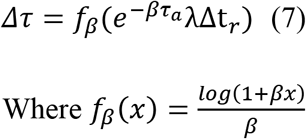

In Eqns. 6 and 7, *f*_*β*_(*x*) is the inverse of *g*_*β*_(*x*) and both functions arise from the integration of Eqn. 5. Eqns. 6 and 7 can be used to convert Δt_r_ into Δτ and vice versa. In turn, Δτ can be converted to H (in °C) and vice versa via the normalisation constant (T_K_).

### Estimating H for a new T_a_

Our main hypothesis is that an organism might have a fixed quantity of a particular currency that, when fully expended during heating, will lead to physiological failure (i.e., T_c_). To test this hypothesis, we estimated what H would be when heated from a different T_a_ for an organism if the amount of currency (the Δt_r_) were indeed a fixed quantity for that organism. This prediction takes the form:

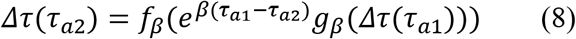

Here, Δτ(τ_a2_) is the unknown heating tolerance starting from τ_a2_ estimated from a known Δτ starting from τ_a1_. That is, we can estimate heating tolerance at a higher temperature (τ_a2_) from heating tolerance at a lower temperature (τ_a1_) (Eqn. 8), and vice versa (Eqn. S7). We used Eqn. 8 to estimate what H at one paired T_a_ would be for the other T_a_, thus assuming that rescaled heating duration (Δt_r_) of the heating event was the same for both experimental pairs. We then compared our estimated heating tolerance H to the true (measured) H for the other T_a_. We also used less derived formulas of Eqn. 8 to represent less derived assumptions for prediction (Supplementary Material). If the estimated and measured H are perfectly correlated, then Δt_r_ is indeed a fixed quantity.

### Exploring the effect of varying heating rate

It has long been recognized that heating rate influences thermal tolerance limits of organisms, so we explored whether heating rate influences rescaled heating duration (Fig. 4A). To do this, we considered a second dataset of T_c_ from 5 phyla (Morley *et al*. 2016) where 37 species had heating tolerance (H) measured under multiple different constant heating rates but starting from the same T_a_ (Supplementary Material). We started with a null hypothesis that rescaled heating duration Δt_r_ does not depend on heating rate λ, so that Δt_r_(λ) = Δt_r_(0). From Eqns. 6 and 7 we have that for any heating rate 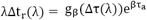 and 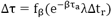). Therefore, under the null hypothesis, Δτ should follow a logarithmic, temperature-dependent relationship with normalised heating rate λ within the observed thermal tolerance and heating rate:

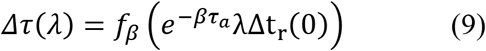

Using the slowest heating rate for each species, we estimated heating tolerance H at all faster heating rates versus observed H under the slowest heating rate and compared with a 1:1 line expected by the null hypothesis. If the null hypothesis is not supported, the points should fall below the identity line. We further visualized the effect of heating rate on rescaled heating duration Δt_r_ by plotting the ratio of rescaled heating duration Δt_r_ of a paired observation with the corresponding ratio for paired heating rate observations λ, within a species. The relationship between the ratio of rescaled heating duration Δt_r_ and the ratio of heating rate λ in the form of an allometric scaling equation Δt_r_(λ)∼cλ^−γ^ is a simple phenomenological expression of how λ influences rescaled heating duration Δt_r_. Parameter c depends only on the experimental protocol and the scaling exponent (γ). A scaling exponent of –1 indicates heating rate does not affect rescaled heating duration Δt_r_.

Indeed, we can insert the expression Δt_r_(λ)∼cλ^−γ^ into Eqn. 7 with heating rate, to estimate Δτ of any species at any T_a_ and under any heating rate λ, given the results from a similar heating assay but performed with a different heating rate and starting temperature T_a_ (Fig. 5C).

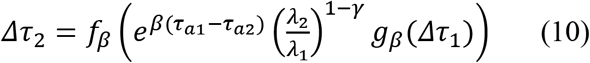

Here, Δτ_2_ is the unknown heating tolerance at heating rate λ_2_, starting from τ_a2_, and estimated from a known Δτ_1_ at heating rate λ_1_, starting from τ_a1_. That is, we can estimate one heating tolerance from another using Eqn. 8 that incorporates the correction for the effect of heating via the exponent γ (Eqn. 10). The same expression can be derived using the inverse 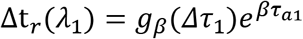 via Eqn. 6. For each species in our datasets that provided data for more than one heating rate, we used Eqn. 10 to estimate every H (as Δτ_2_) using all pairwise combinations of heating rate associated with each known heating tolerance (as Δτ_1_) and a species-specific fitted value of γ.

## Results

Heating tolerance (H) is highly variable within and among species, varying between 3.3 and 37.2°C (mean 15.7°C) among species and acclimation temperatures (Fig. 2A). H was positively but weakly correlated with T_a_ and was notably lower for the higher T_a_ of each species pair (Fig. 2A). The consistently and markedly lower H for higher T_a_ shows ectotherms only elevate their T_c_ a fraction of the corresponding increase in T_a_, which is a well-known pattern (acclimation response ratios tend to be << 1) (Rohr *et al*. 2018). However, rescaling these data to account for the Universal Temperature Dependence of biological rates shows rescaled heating duration, Δt_r_, is strikingly conserved within a species acclimated to very different temperatures T_a_, with all data clustering along the identity line when comparing Δt_r_ for heating assays starting from lower versus higher T_a_ (Fig. 2B).

**Figure 2.**
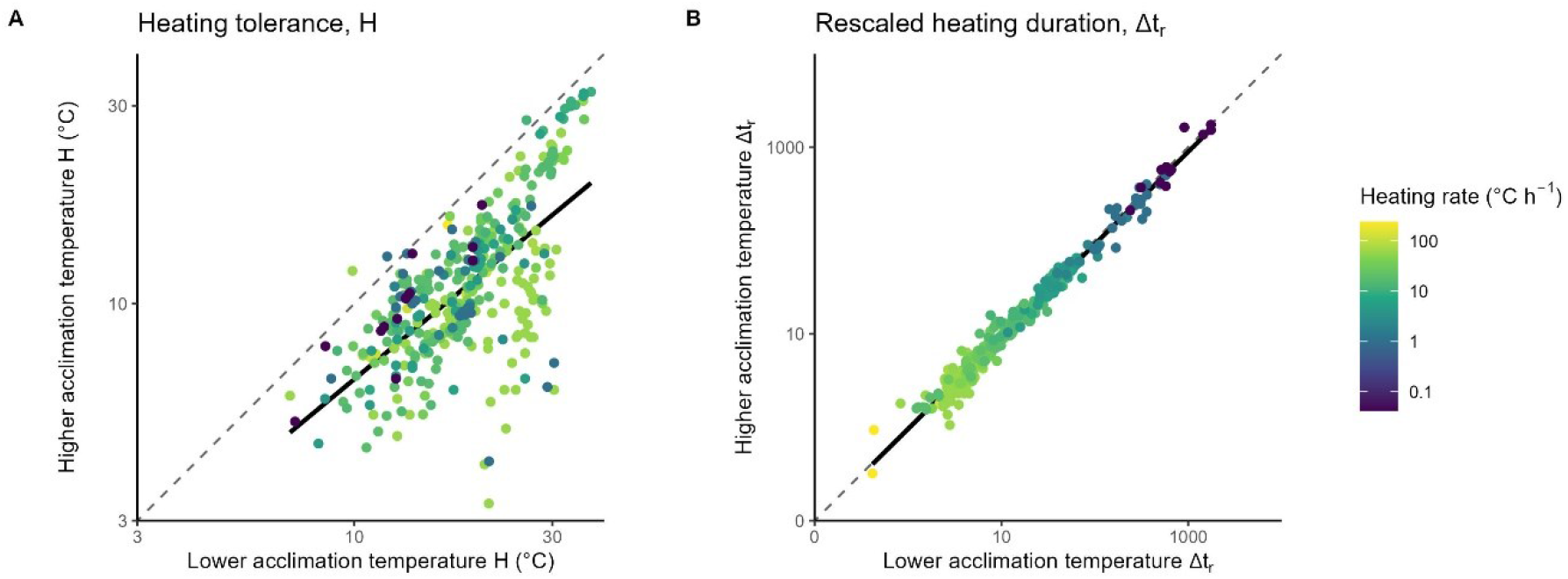
A) Heating tolerance (data points, H) are lower for higher acclimation temperatures, and highly variable across species and heating assays. B) Rescaled heating durations (Δt_r_) corrected for non-linear biological rates are highly conserved at any acclimation temperature, T_a_, across species. Points should fall along the identity line (dashed line) if heating tolerances or rescaled heating durations are the same at both acclimation temperatures and compared with the solid line of an ordinary least-squares regression of the observations. Heating rates (colours) are constant and identical for paired observations. Axes are Log_10_ transformed.

Accordingly, estimating a species’ T_c_ for any new T_a_, if T_c_ is already known for just one other T_a_ (and heating rate is the same), produces remarkably accurate estimates of T_c_ and therefore heating tolerance H (estimated H closely matches observed H; estimating high to low slope = 1.07, R^2^ = 93%; estimating low to high slope = 0.91, R^2^ = 91%, Fig. 3). We found the exponential nature of the scaling along the whole temperature gradient is key: less derived formulations over-or under-estimated Δτ (Figs. S3 & S4). We emphasize that the predictions in Fig. 3 are not formulated by incorporating any phenomenological expression of how T_c_ varies with any other factor (e.g., the *z* parameter, or measuring T_c_ at more than one T_a_); they are made purely by measuring T_c_ for just one T_a_ and by incorporating the scaling of the Universal Temperature Dependence of biological rates (∼ e^−0.6/kT^).

**Figure 3.**
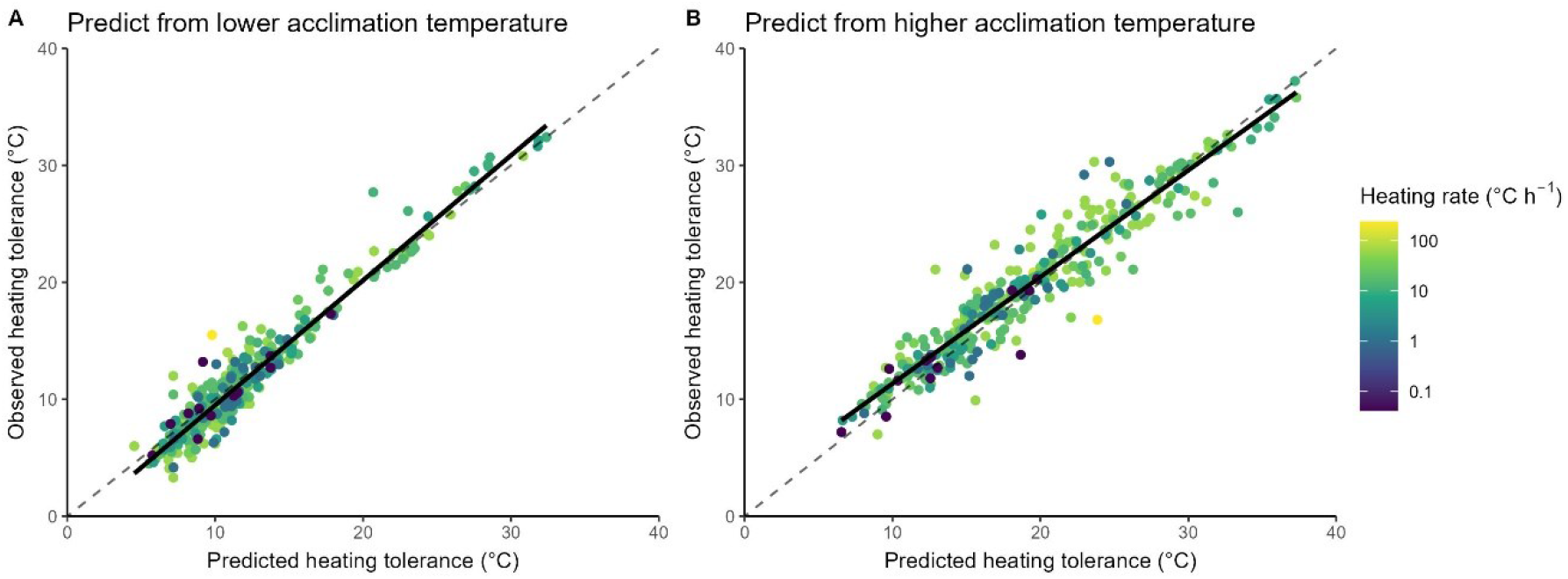
Observed vs predicted heating tolerance (H). A) Predicted heating tolerances at the higher acclimation temperature were calculated from measured rescaled heating duration of the lower temperature heating tolerance and validated against empirically observed heating tolerance at the higher acclimation temperature. B) Predicted heating tolerances at the lower acclimation temperature were calculated from empirically derived rescaled heating duration of the higher temperature heating tolerance and validated against empirically observed heating tolerance at the lower acclimation temperature. Points should fall along the identity line (dashed line) if predictions match observed heating tolerances and compared with the solid line of an ordinary least-squares regression. Heating rates (colours) are constant and identical for paired observations.

Rescaled heating duration Δt_r_ exhibits a strong dependency on heating rate within- and between species, with faster heating reducing the Δt_r_ estimated during heating up to the upper thermal tolerance limit (Fig. 5A; data fall further below the 1:1 line toward larger x-values). This influence of heating rate is also observed in the relationship between heating rate and rescaled heating duration, which is well-described by a power function of form Δt_r_(λ)∼cλ^−γ^ with an interspecific scaling exponent of γ∼0.766, albeit with interspecific variation (Fig. 5B; individually coloured lines). Species-specific γ ranges between –0.06 to –1.05. By incorporating this phenomenological expression (a scaling exponent γ∼0.766) produces good approximations between estimated heating tolerance H (rescaled from Δτ_2_) and observed values of H for all pairwise combinations of heating rate associated with each value of Δτ_1_ (Fig. 5C; data for estimated versus measured H clustered along the identity line). Since Δt_r_ appears to be an approximately fixed property of all species in our dataset for a given heating rate (i.e., no data fall very far from the identity lines in Fig. 2B), and is computed by just a single relationship between temperature and biological rates (the Universal Temperature Dependence), we can incorporate our phenomenological expression that describes how Δt_r_ varies with heating rate to readily generalize how heating tolerance of a hypothetical organism will vary with any T_a_ (Fig. 4B). Finally, by assuming one given quantity of Δt_r_, we can also estimate T_c_ and time to attain T_c_ of a generalised, hypothetical organism for a given acclimation temperature and heating rate (Fig. 4C).

**Figure 4.**
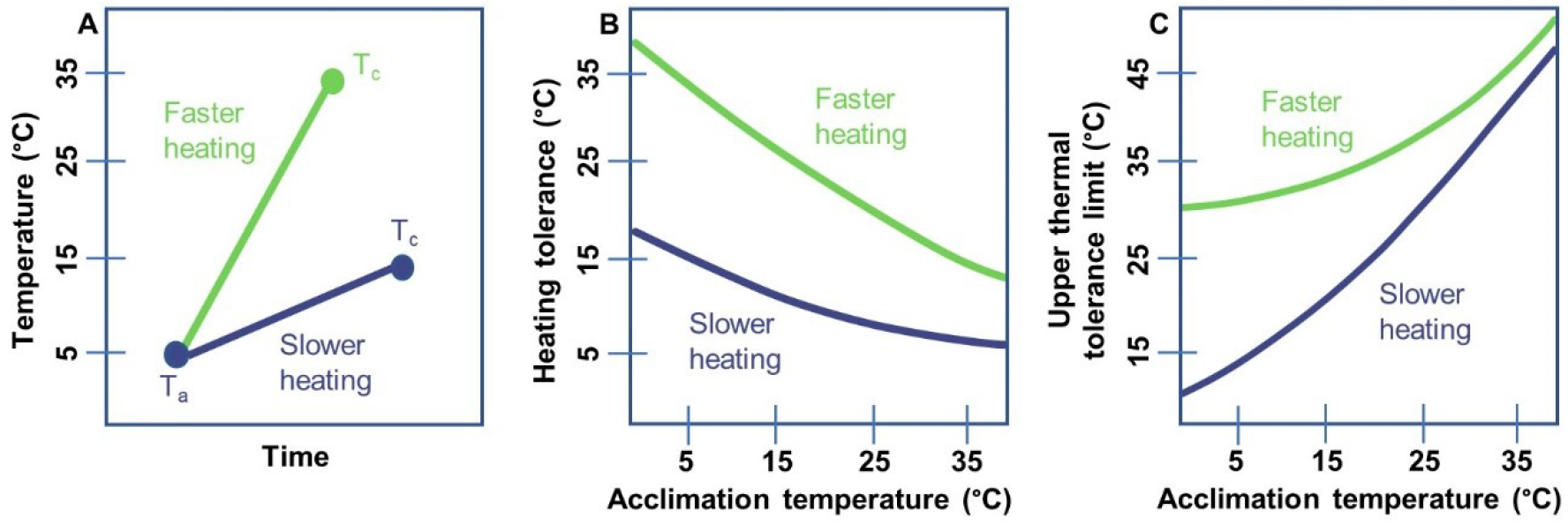
A) For two thermal tolerance assays starting at the same acclimation temperature (T_a_), an assay with a slower heating rate (blue) will have a lower heating tolerance and a longer time (Δt) to reach T_c_ than an assay with a faster heating rate (green). B) Expected heating tolerance (H) decreases with increasing acclimation temperature (T_a_) and is greater at faster heating rates (green) than slower heating rates (blue). C) Expected upper thermal tolerance (T_c_) increases with increasing acclimation temperature (T_a_) and is greater at faster heating rates (green) than slower heating rates (blue). Median values of heating rate and upper thermal tolerance limits at a lower or higher acclimation temperature, as well as a value of 0.766 for γ from the data were used to calculate H and T_c_ (Eqn. 10).

**Figure 5.**
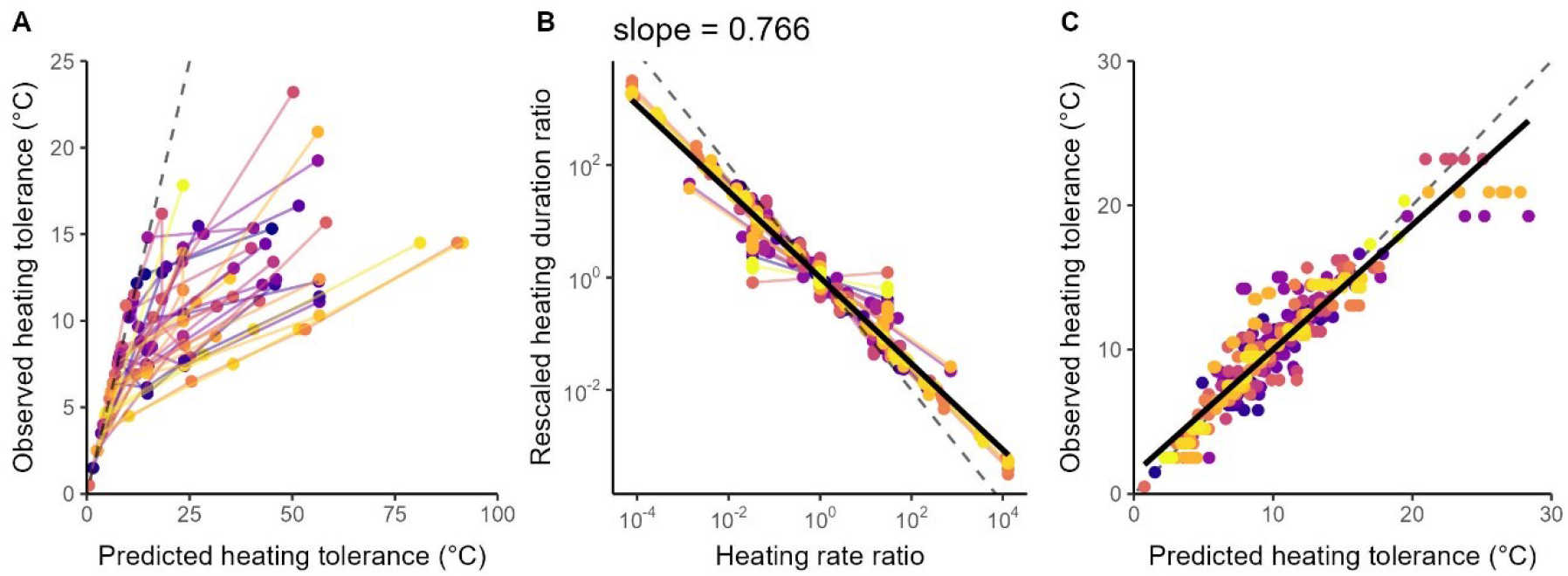
Effect of heating rate on rescaled heating duration. A) Observed heating tolerance (H) against heating tolerance estimated under different heating rates for each species (colours, n = 37). Observations should fall along a 1:1 line (dashed line) if heating rate has no discernible effect on rescaled heating duration (the null hypothesis). Departures from the 1:1 line indicate heating rate reduces rescaled heating duration. B) Ratio of heating rates vs the ratio of rescaled heating durations for all pairs of experiments within a species. Data points represent observations for each species and coloured lines represent ordinary least-squares regressions for each species (n = 37). The solid black line is an ordinary least-squares regression of pooled observations with a slope coefficient of –0.766. Points should fall along the dashed 1:1 line if rescaled heating duration does not depend on heating rate. C) Heating tolerance (H) estimated from observations measured under different heating rates and corrected for the effect of heating rate on rescaled heating durations using species-specific power scaling exponents. The dashed line represents the 1:1 line and the solid line an ordinary least-squares regression of pooled observations.

## Discussion

It is well known that survival probability is determined by the magnitude and duration of thermal stress. Non-linear relationships between temperature and survival time were described more than a century ago (Bigelow 1921), and are now incorporated into concepts of thermal tolerance ‘landscapes’ (Rezende *et al*. 2014). So strong is the temperature-time interaction that quantifying it for a species can subsequently allow accurate predictions of survival times at other static high temperatures, or critical temperatures under different dynamic heating rates (Santos, Castañeda & Rezende 2011; Rezende *et al*. 2014; Kingsolver & Umbanhowar 2018; Jørgensen, Malte & Overgaard 2019; Jørgensen *et al*. 2021). Implicit in this work (and sometimes explicit (Santos, Castañeda & Rezende 2011)) is that thermal limits of an organism have a dependency with the scaling of biological rates with temperature, in part because there exists a temperature-time interaction at all, and in part because of the exponential shape of temperature-survival curves. In this study, we extend these seminal studies and show that the scaling of the Universal Temperature Dependence of metabolism very-closely explains the magnitude that ectothermic animals adjust their critical temperature limits. Importantly, our work requires no phenomenological *a priori* information about how survival times or critical temperatures vary for an organism across any different conditions; if just one T_c_ measurement is known from any T_a_, then T_c_ can be estimated for any new T_a_ with remarkable accuracy (for a given heating rate at least), simply by accounting for the single curve that describes the universal scaling of metabolic rate with temperature (Brown *et al*. 2004). In doing so, our study helps reconcile the fields of physiological thermal tolerance and the metabolic theory of ecology.

Notwithstanding the unambiguous dependency of time on thermal limits (repeatedly observed both in temperature-survival time curves and in relationships between T_c_ and heating rate), thermal tolerance limits are often considered disconnected from the scaling of biological rates with temperature, with the former reflecting structural phenomena (e.g. membrane fluidity)(Bowler 2018), and the latter reaction kinetics (Brown *et al*. 2004). However, structural and reaction rate processes being different mechanisms does not necessarily mean the two are not linked. For example, Bowler (2018) proposed that increases in membrane fluidity are likely to be the ultimate cause of organism death at high temperatures, with thermal perturbation of the plasma membrane precipitating a failure of ion pumps and nutrient transport, impairment of mitochondrial function, a breakdown of homeostasis and, eventually, cell death. But between the ultimate cause of membrane fluidity and proximate causes of cell death, a cascade of processes occurs that are manifested as rates (e.g., rate of nutrient transport or leakage of ions), so it could be expected that the onset of whole-organism failure is indeed regulated by the rate at which biological processes proceed, which are themselves temperature-dependent.

We do not know what combination of specific rate-processes regulate an ectotherm’s tolerance limit, but they seem to be general mechanisms because our dataset consisted of a large number of species (n = 316) from seven Phyla collected across all latitudes and all of them conformed to having a relatively fixed Δt_r_ at different acclimation temperatures (Fig. 2B). Our analyses do not assume that damaging processes only start to occur above some temperature threshold as others do (e.g., Jørgensen *et al*. (2021)) – our Δt_r_ quantity starts accumulating as soon as heating begins – so perhaps an increasing and cumulative departure from homeostasis underlies the currency that, when fully expended, leads to organismal failure. The implication of homeostasis (or at least rapid acclimation processes) is supported by the fact that Δt_r_ was conserved for a species at different T_a_ regardless of whether it was acclimated at T_a_ for 1 day or 300 days. While Δt_r_ appears to be an approximately fixed property of a species (at a given heating rate), it varies by orders of magnitude across species (Fig. 2B) and remains highly variable between species even in a dimensionless form without heating rate (Fig. S2). Some of this variation may come from the different experimental protocols and thermal limit metrics used for different taxa in different experiments, or incomplete compensatory responses during acclimation. But even more variation could arise from differences in the total cumulative departure from homeostasis tolerated by various species before particular cells, tissues and organisms no longer function. Adaptation undoubtedly plays a role too but how our predictions translate to adaptive variation in thermal performance curves is unclear as the Universal Temperature Dependence describes a different relationship of biological rates with temperature than metrics of performance with temperature (Sørensen *et al*. 2018).

We do not contend to have identified any mechanistic (physiological) insight beyond the fact that the scaling of biological rates with temperature almost perfectly explains how an organism varies its physiological temperature limit when acclimated to different temperatures (as long as heating rate is the same). We interpret Δt_r_ as a general pattern rather than a homogeneous pattern and departures from the general trend may identify species or variables of interest for future study. The generality of the model could be tested by comparing results produced with an independent dataset, or by expanding our dataset to include absent organisms with relevant data. Further work could explore what underlies interspecific differences in rescaled heating durations, and also identify the precise mechanistic links between metabolic scaling and thermal limits, because our data suggest those links are strong.

We find a strong effect of heating rate on rescaled heating duration Δt_r_ in our data and show that it is consistent and predictable within and among species following a phenomenological expression. The simple phenomenological expression of how λ influences Δt_r_ can be used to describe the effect of heating rate on rescaled heating durations compared with a null hypothesis (Fig. 5A). The emergence of an approximate negative ¾ power law across species (Fig. 5B) indicates a strong and consistent constraint of faster heating on the rescaled heating durations over which heating is tolerated. This relationship explains how temperature-time dynamics mediate the increase in T_c_ reported for higher heating rates through decreasing the biological-rate-corrected duration of the heating event for an organism. That is, organisms heated faster reach their thermal tolerance sooner on relevant scales of biological rate-corrected time while appearing to reach their thermal tolerance at higher temperatures and have lower heating tolerances on the Celsius scale, in accordance with empirical studies (Peck *et al*. 2009). We can predict the time to attain T_c_ under any heating rate for a known Δt_r_ (Figs. 4B & 4C). If Δt_r_ indexes the cumulative magnitude of an organism’s departure from homeostasis, the negative ∼¾ power dependency of γ on Δt_r_ could reflect some mismatch between supply and demand that becomes increasingly severe as the biological system changes at a faster rate. Such a concept is consistent with the ideas that physiological resistance mechanisms are mostly manifested as rate processes and that thermal tolerance limits are an emergent property of hierarchical and cascading failures of physiological systems (Pörtner 2002; Peck *et al*. 2009). The phenomenological scaling coefficient γ contributes significant predictive power both within and among species, thus using species-specific values for γ provides more accurate predictions of estimated T_c_ than using an interspecific value for γ, however, such information is not always readily available for most species. A fruitful avenue of future work could therefore be to explore what factors influence γ so that more-general predictions can be made with fewer prior measurements. It could be informative to test how our approach applies to static temperatures and tolerance times; we can readily predict how tolerance times vary when an organism is immediately placed at any static high temperature from any T_a_ if we assume a single constant Δt_r_. Testing our predictions with lower thermal limits or cooling events will also be informative of the generality of our model.

The ecological relevance of laboratory-derived tolerance data is regularly questioned, especially for rapid, acute heating experiments (Barnes, Peck & Morley 2010; Clark, Sandblom & Jutfelt 2013; Norin, Malte & Clark 2014; Payne *et al*. 2016), because the influence of temperature on organism performance operates *via* different processes across biological scales (Sinclair *et al*. 2016; Rezende & Bozinovic 2019; Iverson *et al*. 2020). The striking consistencies in the maintenance of rescaled heating durations across broad acclimation periods, temperatures, and heating rates suggest the mechanisms underpinning our data are likely relevant to ectotherms undergoing heating events in nature. Indeed, an increasing number of studies showing correlations between acute organismal responses in the laboratory and biogeographical patterns (Payne *et al*. 2016; Payne *et al*. 2018; Deutsch, Penn & Seibel 2020; Payne *et al*. 2021). Accordingly, our study could complement other recent work that explores the influence of different types of heating events, including fluctuating temperatures (Jørgensen *et al*. 2021), with a view to understand the resilience of organisms exposed to natural heating events in the wild (Bertolini & Pastres 2021). Perhaps one of the most consequential findings of our study concerns the fact that organisms heated at a given rate have practically the same heating tolerance regardless of the temperature they are acclimated to, once heating tolerance is rescaled to account for the Universal Temperature Dependence of biological rates. On one hand, we might interpret this result as ectotherms completely re-adjusting their heating tolerance when they are acclimated to a higher temperature. That is, they exhibit complete metabolic compensation that does not effectively change an organisms’ Universal Temperature Dependence. However, another perspective could be that organisms do not adjust their heating tolerance at all; it is a rigidly fixed property of a species that is unchanged regardless of the temperature they are acclimated to. Either way, our results build on other studies and provide a strong quantitative explanation for why ectotherm thermal tolerance limits vary in the observed manner across macroecological scales, including why they appear to increase only marginally with latitude (Sunday, Bates & Dulvy 2011; Araújo *et al*. 2013) and in acclimation experiments (Gunderson & Stillman 2015; Morley *et al*. 2019b). We contend that those observed patterns do not reflect evolutionary conservatism of thermal tolerance limits, incomplete compensatory metabolic responses, or rigid limits of protein function; they simply reflect a link between thermal tolerance and biological rates, which increase exponentially with temperature (Payne & Smith 2017).

## Acknowledgments

This work was supported by Science Foundation Ireland 18/SIRG/5549 to NLP. ALJ and JFA were funded by an Irish Research Council Laureate Award IRCLA/2017/186. JFA was supported by the “Laboratoires d’Excellences (LABEX)” TULIP (ANR-10-LABX-41). The authors thank Enrico Rezende for helpful discussion and suggestions for improvement. We acknowledge all the animals that provided experimental data to improve our understanding.

## References

Araújo, M.B., Ferri-Yanez, F., Bozinovic, F., Marquet, P.A., Valladares, F. & Chown, S.L. (2013) Heat freezes niche evolution. Ecology Letters, 16, 1206–1219.

Barnes, D.K.A., Peck, L.S. & Morley, S.A. (2010) Ecological relevance of laboratory determined temperature limits: colonization potential, biogeography and resilience of Antarctic invertebrates to environmental change. Global Change Biology, 16, 3164–3169.

Bates, A.E. & Morley, S.A. (2020) Interpreting empirical estimates of experimentally derived physiological and biological thermal limits in ectotherms. Canadian Journal of Zoology, 98, 237–244.

Bertolini, C. & Pastres, R. (2021) Tolerance landscapes can be used to predict species-specific responses to climate change beyond the marine heatwave concept: Using tolerance landscape models for an ecologically meaningful classification of extreme climate events. Estuarine, Coastal and Shelf Science, 252, 107284.

Bigelow, W. (1921) The logarithmic nature of thermal death time curves. The Journal of Infectious Diseases, 528–536.

Bowler, K. (2018) Heat death in poikilotherms: Is there a common cause? Journal of thermal biology, 76, 77–79.

Brown, J.H., Gillooly, J.F., Allen, A.P., Savage, V.M. & West, G.B. (2004) Toward a Metabolic Theory of Ecology. Ecology, 85, 1771–1789.

Chung, D.J. & Schulte, P.M. (2020) Mitochondria and the thermal limits of ectotherms. Journal of Experimental Biology, 223, jeb227801.

Clark, T.D., Sandblom, E. & Jutfelt, F. (2013) Aerobic scope measurements of fishes in an era of climate change: respirometry, relevance and recommendations. Journal of Experimental Biology, 216, 2771–2782.

Deutsch, C., Penn, J.L. & Seibel, B. (2020) Metabolic trait diversity shapes marine biogeography. Nature, 585, 557–562.

Donelson, J.M., Sunday, J.M., Figueira, W.F., Gaitán-Espitia, J.D., Hobday, A.J., Johnson, C.R., Leis, J.M., Ling, S.D., Marshall, D., Pandolfi, J.M., Pecl, G., Rodgers, G.G., Booth, D.J. & Munday, P.L. (2019) Understanding interactions between plasticity, adaptation and range shifts in response to marine environmental change. Philosophical Transactions of the Royal Society B: Biological Sciences, 374, 20180186.

Fredston-Hermann, A., Selden, R., Pinsky, M., Gaines, S.D. & Halpern, B.S. (2020) Cold range edges of marine fishes track climate change better than warm edges. Global Change Biology, 26, 2908–2922.

Gunderson, A.R., Dillon, M.E. & Stillman, J.H. (2017) Estimating the benefits of plasticity in ectotherm heat tolerance under natural thermal variability. Functional Ecology, 31, 1529–1539.

Gunderson, A.R. & Stillman, J.H. (2015) Plasticity in thermal tolerance has limited potential to buffer ectotherms from global warming. Proceedings: Biological Sciences, 282, 20150401.

Hoffmann, A.A., Chown, S.L. & Clusella-Trullas, S. (2013) Upper thermal limits in terrestrial ectotherms: how constrained are they? Functional Ecology, 27, 934–949.

Iverson, E.N.K., Nix, R., Abebe, A. & Havird, J.C. (2020) Thermal Responses Differ across Levels of Biological Organization. Integrative and Comparative Biology, 60, 361–374.

Jørgensen, L.B., Malte, H., Ørsted, M., Klahn, N.A. & Overgaard, J. (2021) A unifying model to estimate thermal tolerance limits in ectotherms across static, dynamic and fluctuating exposures to thermal stress. Scientific Reports, 11, 12840.

Jørgensen, L.B., Malte, H. & Overgaard, J. (2019) How to assess Drosophila heat tolerance: Unifying static and dynamic tolerance assays to predict heat distribution limits. Functional Ecology, 33, 629–642.

Kingsolver, J.G. & Umbanhowar, J. (2018) The analysis and interpretation of critical temperatures. The Journal of Experimental Biology, 221.

Morley, S., Peck, L., Sunday, J., Heiser, S. & Bates, A. (2019a) Physiological acclimation and persistence of ectothermic species under extreme heat events. Global Ecology and Biogeography.

Morley, S.A., Bates, A.E., Lamare, M., Richard, J., Nguyen, K.D., Brown, J. & Peck, L.S. (2016) Rates of warming and the global sensitivity of shallow water marine invertebrates to elevated temperature. Journal of the Marine Biological Association of the United Kingdom, 96, 159–165.

Morley, S.A., Peck, L.S., Sunday, J., Heiser, S. & Bates, A.E. (2018) Acclimation potential of global ectothermic species, collated from literature, 1960 to 2015. Natural Environment Research Council, Cambridge, UK, Polar Data Centre.

Morley, S.A., Peck, L.S., Sunday, J.M., Heiser, S. & Bates, A.E. (2019b) Physiological acclimation and persistence of ectothermic species under extreme heat events. Global Ecology and Biogeography, 28, 1018–1037.

Norin, T., Malte, H. & Clark, T.D. (2014) Aerobic scope does not predict the performance of a tropical eurythermal fish at elevated temperatures. Journal of Experimental Biology, 217, 244–251.

Payne, N.L., Meyer, C.G., Smith, J.A., Houghton, J.D.R., Barnett, A., Holmes, B.J., Nakamura, I., Papastamatiou, Y.P., Royer, M.A., Coffey, D.M., Anderson, J.M., Hutchinson, M.R., Sato, K. & Halsey, L.G. (2018) Combining abundance and performance data reveals how temperature regulates coastal occurrences and activity of a roaming apex predator. Global Change Biology, 24, 1884–1893.

Payne, N.L., Morley, S.A., Halsey, L.G., Smith, J.A., Stuart-Smith, R., Waldock, C. & Bates, A.E. (2021) Fish heating tolerance scales similarly across individual physiology and populations. Communications Biology, 4, 264.

Payne, N.L. & Smith, J.A. (2017) An alternative explanation for global trends in thermal tolerance. Ecology Letters, 20, 70–77.

Payne, N.L., Smith, J.A., van der Meulen, D.E., Taylor, M.D., Watanabe, Y.Y., Takahashi, A., Marzullo, T.A., Gray, C.A., Cadiou, G. & Suthers, I.M. (2016) Temperature dependence of fish performance in the wild: links with species biogeography and physiological thermal tolerance. Functional Ecology, 30, 903–912.

Peck, L.S., Clark, M.S., Morley, S.A., Massey, A. & Rossetti, H. (2009) Animal temperature limits and ecological relevance: effects of size, activity and rates of change. Functional Ecology, 23, 248–256.

Pörtner, H.O. (2002) Climate variations and the physiological basis of temperature dependent biogeography: systemic to molecular hierarchy of thermal tolerance in animals. Comparative Biochemistry and Physiology Part A: Molecular & Integrative Physiology, 132, 739–761.

Rezende, E. L. & Bozinovic, F. (2019) Thermal performance across levels of biological organization. Philosophical Transactions of the Royal Society B: Biological Sciences, 374, 20180549.

Rezende, E.L., Castañeda, L.E., Santos, M. & Fox, C. (2014) Tolerance landscapes in thermal ecology. Functional Ecology, 28, 799–809.

Rohr, J.R., Civitello, D.J., Cohen, J.M., Roznik, E.A., Sinervo, B. & Dell, A.I. (2018) The complex drivers of thermal acclimation and breadth in ectotherms. Ecology Letters, 21, 1425–1439.

Salinas, S., Irvine, S.E., Schertzing, C.L., Golden, S.Q. & Munch, S.B. (2019) Trait variation in extreme thermal environments under constant and fluctuating temperatures. Philosophical Transactions of the Royal Society B: Biological Sciences, 374, 20180177.

Santos, M., Castañeda, L.E. & Rezende, E.L. (2011) Making sense of heat tolerance estimates in ectotherms: lessons from Drosophila. Functional Ecology, 25, 1169–1180.

Sinclair, B.J., Marshall, K.E., Sewell, M.A., Levesque, D.L., Willett, C.S., Slotsbo, S., Dong, Y., Harley, C.D., Marshall, D.J., Helmuth, B.S. & Huey, R.B. (2016) Can we predict ectotherm responses to climate change using thermal performance curves and body temperatures? Ecology Letters, 19, 1372–1385.

Sørensen, J.G., White Craig, R., Duffy Grant, A. & Chown Steven, L. (2018) A widespread thermodynamic effect, but maintenance of biological rates through space across life’s major domains. Proceedings of the Royal Society B: Biological Sciences, 285, 20181775.

Sunday, J.M., Bates, A.E. & Dulvy, N.K. (2011) Global analysis of thermal tolerance and latitude in ectotherms. Proceedings of the Royal Society B: Biological Sciences, 278, 1823–1830.

Terblanche, J.S. & Hoffmann, A.A. (2020) Validating measurements of acclimation for climate change adaptation. Current Opinion in Insect Science, 41, 7–16.

van Heerwaarden, B. & Kellermann, V. (2020) Does Plasticity Trade Off With Basal Heat Tolerance? Trends in Ecology & Evolution, 35, 874–885.

