## Supplementary Material for "Heating tolerance of ectotherms is explained by temperature’s non-linear influence on biological rates"

### Supplementary Material: Heating tolerance of an ectotherm does not vary when temperature's influence on biological rates is accounted for

Jacinta D. Kong[1†], Jean-Francois Arnoldi[2†], Andrew L. Jackson[1], Amanda E. Bates[3], Simon A. Morley[4], James A. Smith[5], & Nicholas L. Payne[1†*]

[1]Trinity College Dublin; Dublin, Ireland.

[2]Centre National de la Recherche Scientifique; Station d'Ecologie Théorique et Expérimentale de Moulis, France.

[3]University of Victoria; Victoria, Canada.

[4]British Antarctic Survey; Cambridge, UK.

[5]University of California Santa Cruz; Santa Cruz, USA.

[†]authors contributed equally.

**Email:**

**Corresponding Author:**

*Nicholas L. Payne.

Department of Zoology

Trinity College Dublin, The University of Dublin,

Dublin Ireland

**Other supplementary materials for this manuscript include the following:**

Dataset S1. Upper critical temperatures of aquatic ectotherms measured under various heating rates, starting from the same acclimation temperature.

Supplementary Code. R script to import and process data, and reproduce results and figures.

#### Heating tolerance data

We reanalyzed a dataset on thermal tolerances of diverse ectotherms containing Acclimation Response Ratios (ARR), acclimation temperatures ($T_a$) and upper thermal tolerance limits ($T_c$) compiled from the literature between 1960 – 2015 [1, 2]. Ectotherms were acclimated for between 1 and 300 days to a lower temperature or a higher temperature for a paired observation for each species. Animals were then heated under a constant ramping temperature from $T_a$ until the point of organismal collapse, which we refer to here as their upper thermal tolerance limit, $T_c$. Thermal tolerance limits were assayed for both acclimation temperatures in paired observations for a species. Within paired observations, each species was measured at the same constant heating rate, but heating rates varied among paired observations representing independent species or heating assays. Data were only included if each species had $T_c$ measured for two different starting $T_a$ of on average 10°C apart, allowing us to compare heating tolerances both within and among species. The dataset contained 369 observations of 316 species from 15 Classes and seven Phyla (Chordata, Arthropoda, Mollusca, Platyhelminthes, Echinodermata, Brachiopoda and Cnidaria) spanning northern and southern hemispheres, polar to tropical latitudes, and freshwater, marine and terrestrial habitats. The $T_c$ of the lower acclimation temperature was not originally reported in [1]. We derive the $T_c$ of the lower acclimation temperature from ARR:

$$T_{c\,lower} = T_{c\,higher} - \left(ARR \times \left(T_{a\,higher} - T_{a\,lower}\right)\right)$$

#### Rescaling heating tolerance

##### Data requirements

Rescaling heating tolerances requires an assay of thermal limits using a ramping protocol with a constant heating rate. Required variables are upper thermal tolerance limit ($T_c$), acclimation temperature or the starting temperature of the ramping assay ($T_a$), and heating rate of the ramping assay (heating rate). These variables are used to calculate heating tolerance and elapsed duration (heating duration) of the ramping assay. Acclimation and starting temperature are used interchangeably as acclimation temperature is the starting temperature of the heating assay in our data. Table S1 lists the main mathematical notation used. The calculations can be generalised to any heating event with a constant heating rate or a known duration of time at a constant temperature. Our model does not formally apply to other protocols of assaying thermal tolerance limits or to lower thermal tolerance limits but these variables present avenues for future studies. Rescaling heating duration and heating tolerance is dependent on three assumptions: 1) Heating tolerance is arbitrarily defined with respect to 0°C, 2) heating rate is constant, and 3) the Universal Temperature Dependence (Arrhenius equation) is not affected by the physiological state of the organism or the experimental conditions (i.e., activation energy E does not change).

#### Physiological rates scale non-linearly with temperature via the Universal Temperature Dependence

As we can expect biological processes occurring during heating to scale with temperature, we can also expect the durations of theses biological processes at one temperature, Δt(T), to scale with temperature. Following the Universal Temperature Dependence, we can describe the relationship between heating durations starting from two temperatures ($T_{a1}$ and $T_{a2}$) as:

$$\frac{\Delta t(T_{a1})}{\Delta t(T_{a2})} \approx e^{\frac{E}{kT_{a1}} - \frac{E}{kT_{a2}}} \qquad \text{(S1)}$$

Here, Δt are elapsed durations of the heating assay starting at temperature $T_a$, E is the activation energy of the Universal Temperature Dependence (0.6eV) and k is Boltzmann's constant. Following Equation S1, we can use a reference temperature to normalize heating durations and effectively derive an expression that is independent of temperature but retains dimensions of time. We can use this normalized temperature scale (τ) to rescale by the non-linear effect of acclimation temperature. We use 0°C as the reference temperature and define a normalization constant ($T_K$) with a value of 273.15 (i.e. 0°C in Kelvin) for centring temperatures (T) on the Celsius scale.

$$\tau = \frac{T}{T_K}$$

Here, T are temperatures on the Celsius scale and τ are normalised temperatures that are centred around the normalisation constant $T_K$. Normalised temperature τ is unitless and can be converted back into degree Celsius by $T = \tau \times T_K$.

Substituting the normalization constant $T_K$ as $T_{a1}$ and temperature T (in °C) as $T_{a2}$ in Equation S1, we can derive an initial expression that rescales heating durations accounting for acclimation temperature:

$$\Delta t_r = e^{\frac{E}{kT_K}} e^{-\frac{E}{kT}} \Delta t(T) \text{ (S2)}$$

Subscript r denotes variables rescaled by the non-linear temperature dependence of physiological rates ("biological rate-corrected", Table S1). $\Delta t_r$ are rescaled heating durations that correspond to non-rescaled heating duration ($\Delta t$).

Rescaled heating duration can be written as an integral between the start ($t_a$) and end times ($t_c$) of the heating assay, assuming a constant temperature ramp:

$$\Delta t_r = e^{\frac{E}{kT_K}} \int_{t_a}^{t_c} e^{-\frac{E}{kT(t)}} dt \qquad \text{(S3)}$$

We can simplify Equation S3 by using the normalized temperature scale ($\tau$) to remove the dimension of temperature. On a normalized temperature scale ($\tau$), $\frac{E}{kT}$ in Equation S3 becomes:

$$\begin{aligned} \frac{E}{kT} &= \frac{E}{kT_K}\frac{1}{1+\frac{T}{T_K}} \\ &= \frac{E}{kT_K}\frac{1}{1+\tau} \\ &\approx \beta(1-\tau) \end{aligned}$$

Where

$$\beta = \frac{E}{kT_K} \qquad \text{(S4)}$$

Here, β (Equation S4) is a dimensionless constant constructed from the organism-level activation energy E, Boltzmann's constant k, and the reference temperature $T_K$. From this, we can approximate the Universal Temperature Dependence as $e^{-0.6/kT} \approx e^{-\beta}e^{\beta\tau}$ and, because we assume β is a constant with an approximate value of 25.5 when E = 0.6 (Table S1), simplify our approximate of the Universal Temperature Dependence as Equation 3 in the main text.

Equation 5 in the main text for rescaled heating duration ($\Delta t_r$) is the simplified form of Equation S3 using the normalized temperature scale ($\tau$) and Equation S4. Rescaled heating duration can be considered as the effective elapsed duration of the heating assay once the non-linear effect of acclimation temperature has been accounted for.

#### Sensitivity analysis

Intraspecific activation energy (E) of various thermally-dependent traits ranges between 0.2 and 1.2 with a median of ~0.6 eV [3]. Changing activation energy and thus β does not meaningfully change the reported patterns (Figure S1) but does show that best alignment of data to the identity line comes from using 0.6-0.8 eV, which is the range best describing the thermal dependence of biological rates and is within the range for the activation energy of whole-organism metabolic rate.

#### Dimensionless rescaled heating duration

A simple way of visualising rescaled heating duration without its dependency on heating rate is to multiply rescaled heating duration by heating rate to give a dimensionless quantity ($H_r = \lambda \Delta t_r$). This dimensionless form of $\Delta t_r$ is expected to be the same at any acclimation temperature and demonstrates the strong relationship between rescaled heating durations at either acclimation temperature, regardless of heating rate (Figure S2). However, variation from the identity line may reflect the effect of heating rate on $\Delta t_r$ and this dimensionless analysis has limited utility in understanding temperature-rate-time dynamics compared with $\Delta t_r$. $\Delta t_r$ is our main aim of our analysis because it includes the integration of the non-linear temperature dependency of biological rates.

#### Predicting heating tolerance for new conditions from a known assay

The simplest model to predict a new heating tolerance from a new acclimation or starting temperature is to assume that heating tolerances are the same for assays starting from the higher or lower temperature. For example, we can predict the heating tolerance at the higher acclimation temperature, $\Delta\tau(\tau_{a2})$, from the lower acclimation temperature, $\Delta\tau(\tau_{a1})$:

$$\Delta\tau(\tau_{a2}) = \Delta\tau(\tau_{a1}) \quad \text{(S5)}$$

Assuming heating tolerance is the same at both acclimation temperatures overestimates the heating tolerance at the higher temperature (Figure S3). The next simplest model is to correct for the difference in acclimation temperatures by rescaling by the acclimation temperature via $e^{\beta(\tau_{a1}-\tau_{a2})}$:

$$\Delta\tau(\tau_{a2}) = \Delta\tau(\tau_{a1})e^{\beta(\tau_{a1}-\tau_{a2})} \quad \text{(S6)}$$

Using Equation S6 underestimates the heating tolerance at the higher acclimation temperature (Figure S4). The fully derived prediction to estimate heating tolerance at one temperature from another temperature is:

$$\Delta\tau(\tau_{a1}) \approx f_\beta(e^{\beta(\tau_{a2}-\tau_{a1})} g_\beta(\Delta\tau(\tau_{a2}))) \qquad \text{(S7)}$$

In Equation S7, $\Delta\tau(\tau_{a1})$ is the unknown heating tolerance starting from $\tau_{a1}$ estimated from a known $\Delta\tau$ starting from $\tau_{a2}$ to estimate heating tolerance at the lower acclimation temperature from the higher acclimation temperature. Equation S7 is the reversed equivalent of Equation 8 in the main text, which estimates heating tolerance at the higher acclimation temperature from the lower acclimation temperature. Equations S7 and Equation 8 in the main text converts normalised heating tolerance at one starting temperature into rescaled heating duration (via $g_\beta$), corrects for the difference in starting temperature along the non-linear Universal Temperature Dependence (via $e^{\beta\tau}$), and converts heating duration corrected for starting temperature and biological rates into normalised heating tolerance at the other starting temperature (via $f_\beta$).

#### Effect of intraspecific heating rate on rescaled heating durations

To examine the relationship between heating rate and rescaled heating duration further, we used a second dataset (Data S1 [4], 107 observations from 5 phyla and 37 species) containing $T_c$ measured from the same $T_a$ under different heating rates ($\lambda$). We combined these observations with the main ARR dataset to produce a final dataset of 127 observations from 5 phyla and 37 species (Supplementary Code).

From heating assays carried out with different heating rates but the same starting temperature, assuming the null hypothesis is accepted, we can infer $\Delta t_r(0)$ from the heating assay with the slowest heating rate $\lambda_{min}$. From Equation 9 in the main text, we should then have that

$$\Delta\tau(\lambda) = f_\beta(\hat{\lambda}), \text{ where } \hat{\lambda} = \frac{\lambda}{\lambda_{min}} g_\beta(\Delta\tau(\lambda_{min})) \qquad \text{(S8)}$$

Thus, plotting the normalized heating tolerance ($\Delta\tau$) against the transformed variable $f_\beta(\hat{\lambda})$ would show if the null hypothesis is supported. If not points would systematically fall below the 1:1 line (Figure 5A in main text).

The relationship between two heating assays done at two heating rates $\lambda_1$ and $\lambda_2$, regardless of the starting temperature, can be described by

$$\frac{\Delta t_r(\lambda_2)}{\Delta t_r(\lambda_1)} \approx \left(\frac{\lambda_2}{\lambda_1}\right)^{-\gamma} \qquad \text{(S9)}$$

Thus, using Equation S9 and estimates of $\gamma$ from the allometric relationship between the ratio of rescaled heating duration and the ratio of heating rate (Figure 5B in main text), we can deduce $\Delta t_r(\lambda_2)$ we can derive Equation 10 in the main text to predict heating tolerances accounting for the effect of heating rate on rescaled heating duration (Figure 5C in main text).

#### Supplementary Figures and Tables

Table S1 Descriptions of mathematical notations used

| Notation | Description | Units (if applicable) | Value (if constant) |
|---|---|---|---|
| H | Heating tolerance; range of temperatures between $T_a$ and $T_c$ | °C | |
| T | Temperature | °C unless otherwise stated as K | |
| r | Subscript denotes variables that have been rescaled to correct for non-linear biological rate-temperature relationships (the Universal Temperature Dependence) | | |
| a | Subscript denotes variables associated with acclimation or starting temperature | | |
| c | Subscript denotes variables associated with upper thermal tolerance limit | | |
| E | Activation energy of Universal Temperature Dependence | eV | 0.6 |
| $T_K$ | Normalisation constant, Reference temperature for centring | K | 273.15 (0°C) |
| t | Time | Time | |
| τ | Normalized temperature; centred around $T_K$ | Dimensionless | |
| β | Biological rate dependency parameter. Derived from the Universal Thermal Dependence | | Approx. 25.5 with E = 0.6 |
| $g_\beta(x)$ and $f_\beta(x)$ | Parameter functions from integrating the normalised Universal Thermal Dependence | | |
| $\Delta t$ | Duration of heating assay between $t_a$ and $t_c$ | Time | |
| $\Delta t_r$ | Biological rate-corrected duration of heating assay (rescaled heating duration); effective duration of heating assay | Time | |
| $\Delta\tau$ | Normalized heating tolerance between $\tau_a$ and $\tau_c$. Not accounting for heating rate or the Universal Temperature Dependence | Dimensionless | |
| Heating rate | Constant rate of temperature increase in the heating assay | °C $h^{-1}$ | |
| λ | Heating rate on the normalised temperature scale | $Time^{-1}$ | |
| γ | Allometric coefficient for the effect of heating rate on rescaled heating duration $\Delta t_r$ | | Fitted from data |

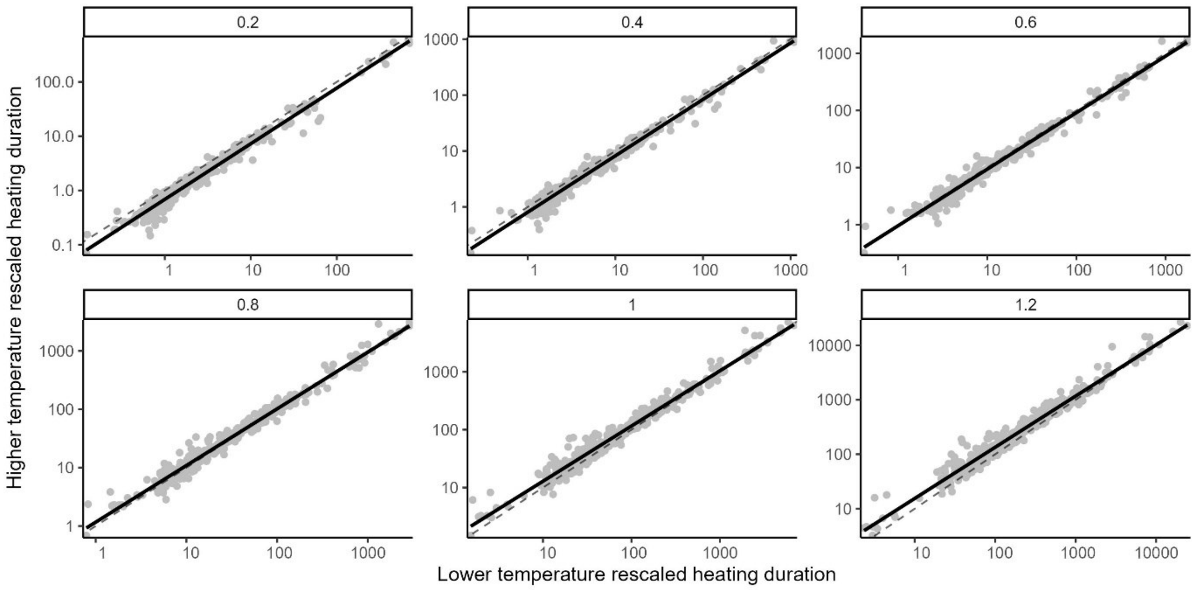


Figure S1. Relationships between heating tolerance expressed as biological rate-corrected heating duration $\Delta t_r$ (rescaled heating duration, points) at two acclimation temperatures are similar across a range of activation energies used (0.2 – 1.2 eV). The dashed identity line represents the expected relationship if rescaled heating duration was the same at both acclimation temperatures and the solid line is an ordinary least-squares regression. Axes are $Log_{10}$ transformed.

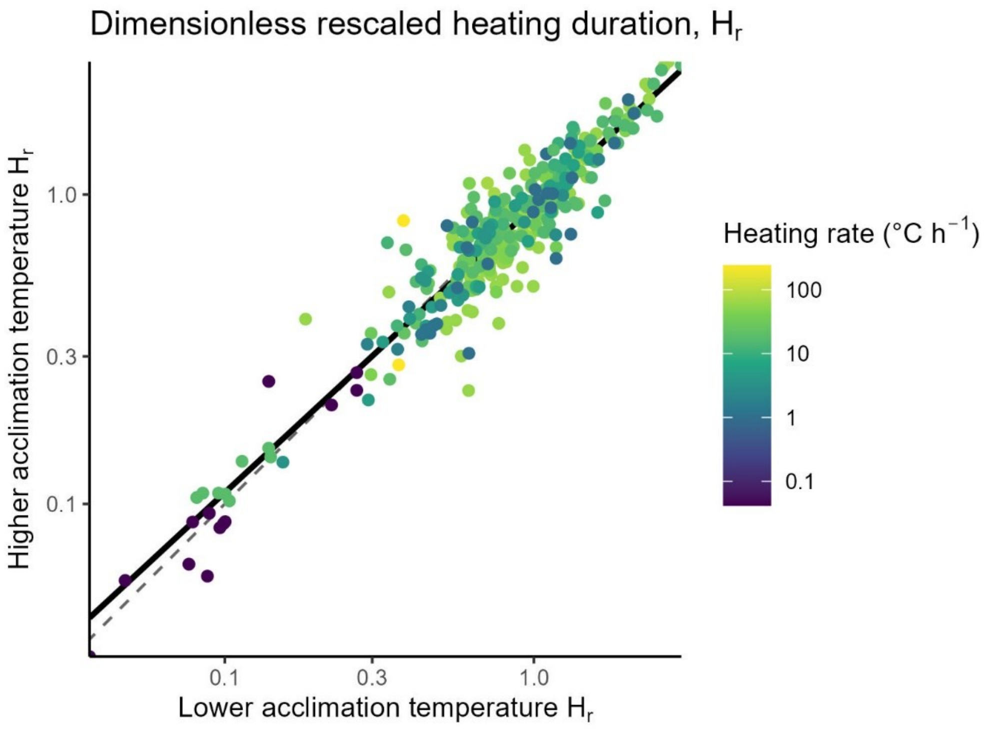


Figure S2. Dimensionless heating duration ($H_r$) corrected for non-linear biological rates and accounting for heating rates are similar between acclimation temperatures, and vary within one order of magnitude across experiments and organisms. Points should fall along the identity line (dashed line) if ectotherms fully compensate their thermal tolerance limits, $T_c$, at higher $T_a$, and compared with the solid line of an ordinary least-squares regression. Heating rates (colours) are constant and identical for paired observations. Axes are $Log_{10}$ transformed.

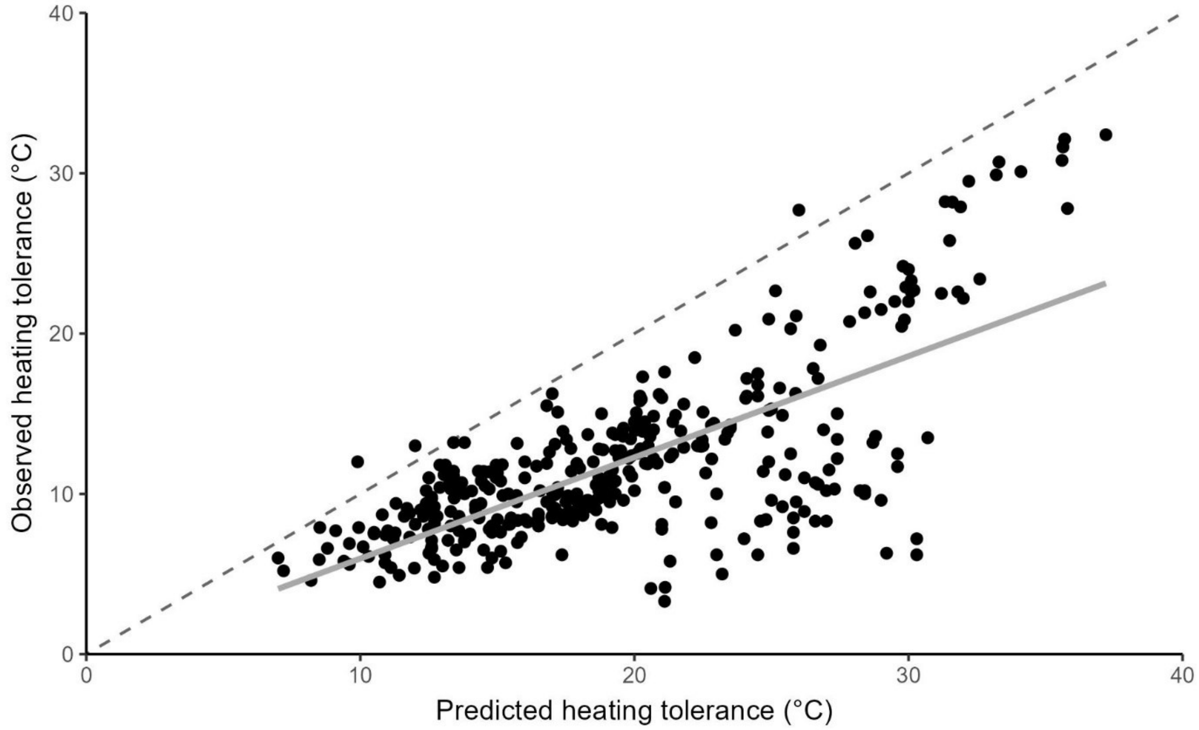


Figure S3. Assuming that the lower acclimation temperature heating tolerance (H, °C) is the same as the higher acclimation temperature heating tolerance (H, °C) overestimates heating tolerance at the higher temperature. Predicting a new heating tolerance is much less accurate then when rescaled heating duration is accounted for (compare with Figure S4). Predictions were made for each species (points). Predictions should fall along a 1:1 identity line (dashed line) if assuming that the heating tolerance of the higher temperature is the same as at the lower temperature is accurate. The solid line is an ordinary least-squares regression.

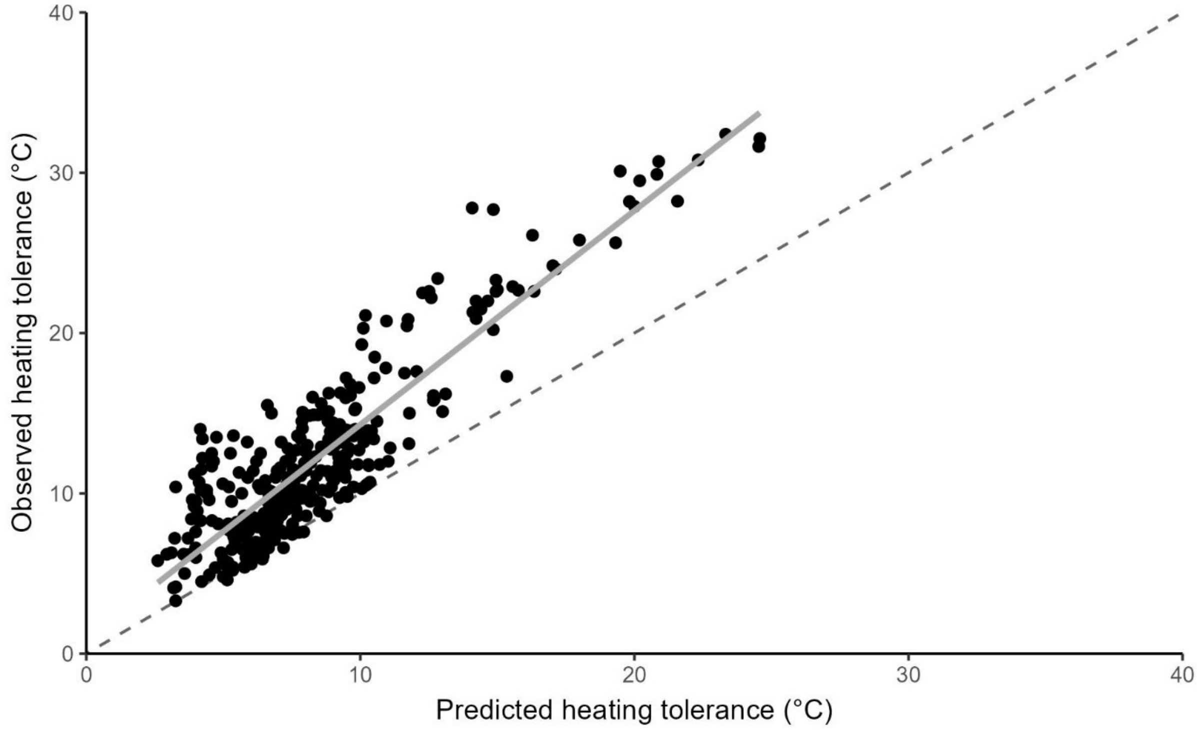


Figure S4. Correcting for the change in acclimation temperature by rescaling the lower acclimation temperature (via $e^{\beta(\tau_{a2}-\tau_{a1})}$) underestimates the heating tolerance (H, °C) at the higher acclimation temperature. As in Figure S3, this shows that failing to account for acclimation temperatures along the Universal Temperature Dependence leads to less accurate predictions of heating tolerance (H, °C). Predictions were made for each species (points). Predictions should fall along a 1:1 identity line (dashed line) if accounting for the difference in acclimation temperature by rescaling the acclimation temperature is adequate. The solid line is an ordinary least-squares regression.

#### References

[1] Morley, S.A., Peck, L.S., Sunday, J.M., Heiser, S. & Bates, A.E. 2019 Physiological acclimation and persistence of ectothermic species under extreme heat events. *Glob Ecol Biogeogr* **28**, 1018-1037. (doi:10.1111/geb.12911).
[2] Morley, S.A., Peck, L.S., Sunday, J., Heiser, S. & Bates, A.E. 2018 Acclimation potential of global ectothermic species, collated from literature, 1960 to 2015. (Polar Data Centre, Natural Environment Research Council, Cambridge, UK.
[3] Dell, A.I., Pawar, S. & Savage, V.M. 2011 Systematic variation in the temperature dependence of physiological and ecological traits. *Proc Natl Acad Sci U S A* **108**, 10591-10596. (doi:10.1073/pnas.1015178108).
[4] Morley, S.A., Bates, A.E., Lamare, M., Richard, J., Nguyen, K.D., Brown, J. & Peck, L.S. 2016 Rates of warming and the global sensitivity of shallow water marine invertebrates to elevated temperature. *J Mar Biol Assoc U K* **96**, 159-165. (doi:10.1017/S0025315414000307).
